# Prestimulus neural variability affects behavioral performances mediated by poststimulus-evoked responses at the intraindividual and interindividual levels

**DOI:** 10.1101/2023.06.26.546352

**Authors:** Zeliang Jiang, Xingwei An, Shuang Liu, Erwei Yin, Ye Yan, Dong Ming

## Abstract

There are significant intra-individual and inter-individual variabilities in audiovisual temporal perception. Previous studies have shown that prestimulus neural variability could reflect behavioral variabilities. We aimed to investigate whether prestimulus neural variability can predict behavioral variability in audiovisual temporal perception. Furthermore, We also explored whether prestimulus neural variability directly influences behavioral responses or indirectly impacts perceptual decisions through post-stimulus-evoked responses. We analyzed the electroencephalography (EEG) data from a paradigm where the twenty-eight human subjects performed a simultaneity judgment (SJ) task in the beep-flash stimulus. The prestimulus weighted permutation entropy (WPE) was the indicator of neural variability in this study. We found that prestimulus frontal WPE could predict the individual’s TBW in auditory- and visual-leading conditions. In addition, increased prestimulus parietal WPE was associated with more asynchronous responses. Prestimulus frontal WPE may be associated with top-down cognitive control, while parietal WPE may be related to bottom-up cortical excitability. Furthermore, poststimulus evoked responses could mediate the relation between prestimulus WPE and the individual’s TBW or perceptual responses. These results suggested that prestimulus WPE was a marker in reflecting intra-individual and inter-individual variabilities in audiovisual temporal perception. Significantly, prestimulus WPE might influence perceptual responses by affecting poststimulus sensory representations.

## 1 Introduction

When we face the same stimulus, our responses to the stimulus show large variability across different individuals, referred to as inter-individual differences ^[1]^. Even when the stimulus is repeatedly represented, our responses differ over trials, referred to as intra-individual differences ^[1]^. These variabilities could be reflected by poststimulus-evoked responses such as ERP ^[2]^, ERSP ^[3]^, and brain network ^[4]^. Some recent studies began to use prestimulus spontaneous neural oscillations ^[3, 5-7]^ to characterize them.

Research has recently revealed that the neural variabilities in the brain, previously considered meaningless noise or measurement error [8, 9], reflect these behavioral variabilities. Neural variability is believed to contain valuable information about brain dynamics, which is not only related to low- and high-level cognition in health and clinical participants ^[8]^ but also underlies the plasticity and adaptability of human behavior ^[9, 10]^. Brain neural variability can take many forms. In studies utilizing electroencephalography (EEG) and magnetoencephalography (MEG), it can be broadly classified into two categories: one is moment-to-moment variability which reflects how much brain activity changes from one moment to the next, that is, the brain activity range and can be quantified by entropy; another is trial-by-trial variability (TTV) which reflects the distributional width of a neural time series and can be measured by the variance or standard deviation of neural signals ^[9, 11, 12]^.

Entropy correlates with local cortical desynchronization ^[13, 14]^. Large local cortical desynchronization will increase entropy and further influence stimulus processing. For instance, using a pitch discrimination task, Waschke et al. (2017) found that increased prestimulus weighted permutation entropy (WPE) in auditory-related channels raised pitch detection rates. More importantly, they rose with age ^[14]^. Of note, in the study of Waschke et al. (2017), as an inter-individual difference marker in aging, WPE is task-free and trait-like. So whether the WPE also predicts task-induced and state-like individual differences remains to be studied. Previous studies showed that the alpha frequency band exhibits a unique role in individuality in high-level moral decision-making ^[15]^, language comprehension ^[16]^, and low-level perceptual judgments ^[17-19]^. Given that WPE is related to the power of low-frequency (1-30 Hz) neural oscillations ^[13, 20]^, we speculated that prestimulus WPE might also predict task-induced and state-like individual differences.

Perceptual decision-making typically involves early sensory representations and later decision-related computations via poststimulus brain response analysis ^[21-23]^. Recently, increasing numbers of studies found prestimulus spontaneous neural oscillations shape behavioral responses by modulating cortical excitability corresponding to perceptual modalities ^[5, 24-26]^ and attentional state ^[27]^. Furthermore, some previous studies attempted to examine the interrelation between spontaneous neural oscillations, task-induced activity, and behavior. However, the results were inconclusive: some studies pointed out that the prestimulus state affects perception mediated by task-induced activity ^[28-30]^, while others found prestimulus state directly affects perception. Although few studies have directly investigated how prestimulus neural variability affects behaviors by the task-induced response ^[29, 31]^, some papers found that sensory encoding and cortical representations quantified by the power or ITC of neural oscillations depend on the degree of neural variability in no-human animals ^[32-34]^ and human subjects ^[13, 14, 35]^. Then, we speculated that prestimulus neural variability shapes behavior mediated by sensory encoding and cortical representations.

This study first aimed to test whether prestimulus neural variability quantified WPE predicts intra- and inter-individual behavioral variations in audiovisual temporal perception. Then, if any, we wanted to further clarify the functional role of prestimulus neural variability on behavioral variations by investigating the interrelation between variability of ongoing brain signals, evoked responses, and behavior. We reanalyzed the EEG data from a previous study acquired while a group of young adults was engaged in an audiovisual simultaneity judgment (SJ) task to find the neural correlates of audiovisual temporal binding. As we know that the temporal binding windows (TBW) measured by the SJ task vary substantially between individuals ^[36-38]^, we were able to study the inter-individual differences. In the EEG experiment, subjects make different judgments when the same stimulus (i.e., the individually specific points of the perceptual uncertainty where the probability of synchronous and asynchronous responses is equal: ∼50%) was repeatedly presented. So we can study the intra-individual variations (see 2.3 task design). Given that audiovisual temporal binding engages different neural pathways depending on the leading sense (audiovisual [AV] or visual-audio [VA]) ^[39, 40]^, we explored the mechanisms of neural variability in AV and VA conditions separately.

## 2 Material and methods

### 2.1 Participants

Twenty-three healthy young adults were recruited in the previous study, and we additionally recruited five participants. There are twenty-eight young adults (9 male; 21-29 years old) finally. We separately focused on all trials regardless of response types in auditory-leading stimulus onset asynchrony (SOA) and visual-leading SOA in inter-individual difference data analysis. So all 28 subjects were included in the inter-individual difference data analysis. However, in intra-individual difference data analysis, we focused on the signal difference between different response types. Given that the probability of synchronous and asynchronous responses is extremely unbalanced for the fifteenth subject in visual-leading SOA and the twentieth subject in auditory-leading SOA, the data were excluded in the final data analysis. There are 27 subjects in the intra-individual difference data analysis (see Table 1). All subjects were right-handed. They all had normal or corrected to normal vision and normal hearing. The study was conducted following the tenets of the Declaration of Helsinki and was approved by the Ethics Committee of Tianjin University.

**Table 1.** The distribution of synchronous judgment ratios in each subject

| Participant number | A50V(%) | V50A(%) |
| --- | --- | --- |
| 1 | 68.00 | 34.00 |
| 2 | 27.54 | 60.00 |
| 3 | 77.36 | 72.78 |
| 4 | 57.59 | 84.18 |
| 5 | 79.81 | 46.23 |
| 6 | 64.42 | 47.09 |
| 7 | 83.72 | 76.98 |
| 8 | 77.40 | 52.17 |
| 9 | 36.06 | 63.90 |
| 10 | 37.38 | 47.32 |
| 11 | 32.14 | 48.57 |
| 12 | 57.14 | 34.78 |
| 13 | 66.43 | 21.74 |
| 14 | 66.67 | 51.45 |
| 15 | 73.57 | 4.380 |
| 16 | 42.14 | 52.90 |
| 17 | 41.43 | 74.10 |
| 18 | 24.29 | 54.35 |
| 19 | 30.94 | 45.26 |
| 20 | 94.29 | 38.57 |
| 21 | 52.55 | 57.97 |
| 22 | 57.86 | 70.00 |
| 23 | 50.71 | 70.59 |
| 24 | 77.14 | 17.99 |
| 25 | 72.99 | 36.36 |
| 26 | 59.29 | 52.52 |
| 27 | 28.57 | 51.80 |
| 28 | 50.71 | 61.15 |

Note. The A50V SOA represented auditory-leading SOA, while V50A SOA represented visual-leading SOA. The conditions with extremely unbalanced judgment ratios in red font were excluded from the intra-difference analysis.

### 2.2 Task design

The task was designed with the Psychophysics Toolbox, version 3.0.11, with MATLAB R2017a (MathWorks). The participants were required to perform two sessions: The first session is a behavioral experiment of 4 blocks, each with 13 stimulus onset asynchronies (SOAs) (± 300, ± 250, ± 200, ± 150, ± 100, ± 50, and 0 ms) for the first ten subjects or 15 SOAs (± 400, ± 300, ± 250, ± 200, ± 150, ± 100, ± 50, and 0 ms; a positive SOA denotes visual-leading presentation) for the last eighteen subjects, and five trials per SOA, aiming at obtaining the width of TBWs (visual leading TBW: RTBW; auditory leading TBW: LTBW), points of perceptual uncertainty (A50V and V50A SOA: the 50% proportion of synchronous responses in auditory or visual leading SOA) and Point of Subjective Simultaneity (PSS: the point with the highest proportion of synchronous reactions). We used two psychometric sigmoid functions to fit the data in behavioral sessions. The second session is an EEG experiment with eight blocks of 56 trials (± 300 ms: 6, PSS:6, A50V:26, V50A:26, the first ten subjects) or seven blocks of 52 trials (± 300 ms: 6, PSS:6, A50V:20, V50A:20, the last eighteen subjects) aimed at obtaining the EEG data of intra-individual variability ^[37]^.

The visual stimuli used were white rings (outer diameter= 10.5°, inner diameter = 8°) and presented on a Philips 236V6Q monitor at a refresh rate 60Hz with 33 ms (2 frames) duration. Auditory stimuli were pure tones (1800 Hz) presented for 33 ms (including 2.5 ms fade-in and 2.5 ms fade-out) via an In-Ear-Monitor (EDIFIER H230P). The relative timing of auditory and visual stimuli was tested to be precise up to about 3 ms using an audio cable, a photodiode, and a SynAmps amplifier system (Neuroscan).

The experimental process was as follows: First, a pair of beep-flash appears following a white central cross lasting about 1500-2000 ms. After the audiovisual stimulus, a white cross of 500 ms again emerged as a response delay. Third, a response interface appeared. At this point, the white cross turned green. The subjects were asked to perform the two-alternative forced-choice (2AFC) simultaneity judgment task. Press the “F” button when the stimuli are synchronous, and press the “J” button when they are asynchronous.

### 2.3 EEG acquisition and preprocessing

The EEG signal was recorded using a 62-channel Neuroscan system with scalp electrodes placed according to the International 10–20 electrode placement standard, with the left mastoid reference and ground between FPz and Fz. The acquisition rate was 1000 Hz. EEG data analysis was performed with the EEGLAB toolbox versions 13.6.5b ^[41]^ and Fieldtrip Toolbox ^[42]^. All the preprocessing steps of EEG data were performed using the EEGLAB toolbox. EEG channels were firstly re-referenced to two linked mastoids and down-sampled to 500 Hz. Then the data were band-pass filtered (1-80 Hz) and notch filtered (around 50 Hz) with a Hamming windowed-sinc finite impulse response zero-phase filter. The ICA algorithm and ADJUST plugin of EEGLAB were used to eliminate artifacts from the data. The clean data were segmented into epoched from -1500 ms before the first stimulus onset to 1500 ms. Epochs containing voltage deviations exceeding ±100 μV were excluded.

### 2.4 EEG analysis

#### 2.4.1 Time-Frequency analysis

Single-trial complex valued time-frequency (TF) representation was obtained using wavelet analysis in FieldTrip. Frequencies ranged from 3 to 30 Hz in steps of 1 Hz, while the width of individual Morelet wavelets was fixed at three cycles. TF data were calculated at time points of -1.5 to 0.3 s peristimulus in a 10-ms bin, as it has been shown that cumulative sensory evidence generally occurs within 300 ms of stimulus onset ^[23]^. The power was first calculated by squaring the modulus of the time-frequency spectrum. Next, baseline-corrected relative power was expressed as a change in percentage (%) relative to average power in the -1.5 to -1 s time window as in entropy-related analysis.

Additionally, we divided the time-frequency spectrum by its magnitude and averaged it across trials to compute the magnitude of the average complex value to get the inter-trial phase coherence (ITC) and evaluated how consistent the oscillatory phase is across an ensemble of trials for inter-indifference analysis. For intra-individual variability analysis, we use the Jackknife method to obtain the phase coherence of a single trial which was referred to as jackknife-ITC (jITC) ^[13, 43]^ by calculating the ITC of all trials but the one in question. The jITC calculated the ITC of other trials except this trial. If the contribution of this trial to all trials is large, then the magnitude of jITC in this trial will become smaller. According to Waschke et al. (2019), we used 1–jITC to represent single-trial ITC comparable to ITC across trials ^[13]^. Finally, the single-trial ITC was logit-transformed for statistical analysis.

#### 2.4.2 Entropy calculation

We computed weighted permutation entropy (WPE) ^[44]^ of EEG signals which quantified moment-to-moment variability in a moving window fashion and were used extensively in neuroscience domains ^[13, 14, 20, 45]^. We used the Matlab wpH.m script written by Waschke et al. (2019) to quantify the time course of single trial WPE [details can be found in Waschke et al. (2019), Fadlallah et al. (2013)] ^[13, 44]^. According to previous research, we use 50 samples moving window (100 ms in this study), 10 sample points (20 ms) in step, and 3 sample points in calculating every Shannon entropy. We used relative WPE to quantify neural variability. So we selected the time interval of -1.5s to -1s as a baseline for baseline correction to get the relative WPE as was done in similar paper ^[14]^.

### 2.5 Statistical Analysis

#### 2.5.1 Behavior statistical analysis

To verify whether our grouping method is reasonable, we used an independent samples t-test to examine whether the difference of individual TBWs between groups was significant.

#### 2.5.2 EEG statistical analysis

We used a nonparametric cluster-based permutation procedure in EEG statistical analysis to assess the statistical significance. Specifically, first, paired-sample/independent samples t-test or Pearson correlation analysis was used to get the raw statistical values (t-values in Fieldtrip). Then, this procedure was performed 1000 times while shuffling the labels across subjects or reactions randomly each time to generate a null distribution and get the null-level statistical values. Finally, all t-values above a threshold corresponding to an uncorrected p-value of 0.05 were formed into clusters by grouping adjacent at least two significant time, frequency, and electrode points in the raw and shuffling statistics. When the raw clusters were higher than the 97.5th percentile or lower than the 2.5th percentile of this null distribution, the clusters were considered to have a significant effect (5% alpha level). In the inter-individual variation analysis, the EEG data were averaged over synchronous and asynchronous responses in the A50V and V50A SOA for LTBWs and RTBWs, respectively. In the intra-individual variation analysis, the EEG data were analyzed between two different reactions in the A50V and V50A SOA condition, respectively.

#### 2.5.3 EEG-behavior correlation/regression and mediation analysis

We used the mediation model to reveal the relationship between prestimulus and poststimulus brain responses and behaviors.

In the analysis of the inter-individual differences, we first used Pearson correlation analysis to find the relationship between prestimulus (WPE) and poststimulus brain response (ITC or power) and between poststimulus brain response and behavior (that is, Brain-Brain and Brain-Behavior model). Next, we extracted the time-frequency and channel points whose distributions of correlation coefficients are consistent in the above two analyses and averaged them. Then, we used the PROCESS macro ^[46]^ in SPSS for mediation analysis (the Brain-Brain-Behavior model). The significance of the path coefficients in the mediation model was assessed with the bootstrap method. The 90% bootstrap confidence intervals (CIs) of the path coefficients were calculated based on 5000 bootstrap samples given the small sample size.

In the intra-individual differences analysis, we tried to find the Brain-Brain-Behavior model at the single-trial level. We first used single-trial estimates of pre- and poststimulus brain activity as well as behavioral response (‘synchronous’ vs. ‘asynchronous’) to fit three models: model 1, prestimulus brain response (WPE) as predictors while behavioral response as dependent variables to fit a logistic regression model; model 2, prestimulus brain response (WPE) as predictors while poststimulus brain response (jITC or power) as dependent variables to fit a linear regression model; model 3, poststimulus brain response (jITC or power) as predictors while behavioral response as dependent variables to fit a logistic regression model. Next, we compared the slopes of each data point across subjects in each model against zero using a one-sample t-test to get data points with significant and consistent differences across subjects in three models. Finally, we extracted consistent data points to perform mediation analyses in SPSS. The 95% bootstrap confidence intervals (CIs) of the path coefficients were calculated in this analysis.

## 3 Results

### 3.1 Behavior Results

The subjects were split into three groups (triple-split) ^[47]^ for the LTBW and RTBW, respectively. We choose the top and bottom third subjects for between-group comparison (Fig. 1a). The inter-group comparison of TBWs showed that the width of TBWs in the two narrow TBW groups was significantly narrower than that in the wide TBW groups [LTBW group: *t*_(17)_ = -5.467, *p* < 0.001, Cohen’s d = -2.512; Narrow_LTBW = 111.608 ± 24.491 ms, Wide_LTBW = 249.214 ± 71.662 ms (mean ± SD); RTBW group: *t*_(17)_ = -6.422, *p* < 0.001, Cohen’s d = -2.951; Narrow_RTBW = 165.173 ±27.844 ms, Wide_RTBW = 264.652 ± 38.178 ms (mean ± SD); Fig. 1b]. The behavioral results suggest that visual and auditory leading TBW groupings effectively differentiate subjects with narrower and broader TBWs.

**Fig. 1.**
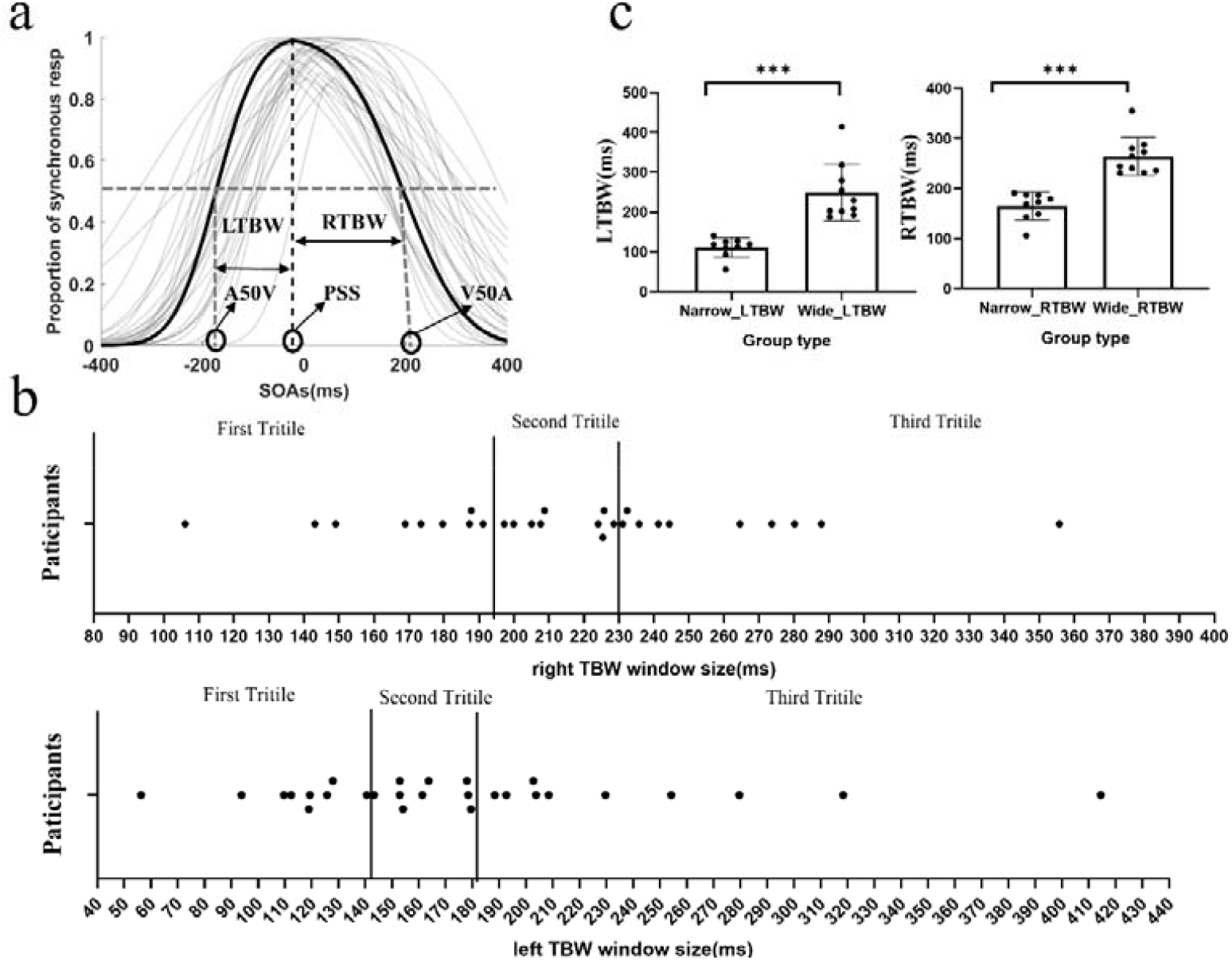
Distribution of subjects’ TBWs and results of the inter-group comparison. (a) The grey curves represent the two fitted sigmoid curves based on each subject’s behavioral data, while the dark black curves represent the group-averaged data. Negative numbers represent auditory-first Stimulus Onset Asynchronies (SOAs), while positive numbers represent visual-first SOAs. (b) The Distribution of subjects’ LTBW and RTBW in the top and bottom subpanels. The participants were divided into three groups by tertiles to maximize between-group differences. Each point represents a subject. We used the first and third tertiles for statistical comparison. (c) The results of the inter-group comparison in the LTBW and RTBW. Narrow_LTBW or Narrow_RTBW represents narrower auditory-leading or visual-leading TBWs, while Wide_LTBW or Wide_RTBW represents wider auditory-leading or visual-leading TBWs. * p < 0.05, ** < 0.01, *** < 0.001.

### 3.2 Prestimulus WPE reflects inter-individual differences in TBW mediated by poststimulus ITC

In the RTBW group, we extracted and computed the mean of prestimulus WPE from -500 to -400 ms before the onset of the visual stimulus. Subsequently, we employed an independent samples t-test (Wide_RTBW contrast Narrow_RTBW) combined with a spatial cluster-based permutation test to analyze the data. We found a significant positive cluster over prefrontal and central sensors (cluster-value = 61.714, *p* = 0.01; fig. 2a), suggesting an increase in prestimulus WPE reflected weaker temporal perception. Further, we examined the relationship between prestimulus WPE and RTBW across subjects using Pearson’s correlations. Positive correlations were significant between the prestimulus WPE and the magnitude of RTBW over prefrontal, central, and parietooccipital sensors (cluster-value = 102.065, p = 0.002; fig. 2b). Given that prestimulus WPE is related to prestimulus low-frequency power ^[13, 14]^, we extracted and averaged the prestimulus WPE over the significant time interval and sensors and entered this into Pearson’s correlation analysis with prestimulus low-frequency power with the spatiotemporal-electrode cluster-based permutation test. Significant negative correlations were found, which were mainly concentrated in the theta band (4 -7 Hz) of -0.5 to -0.4 s over prefrontal and central sensors (cluster-value = -49427, p = 0.018; fig. 2d). We further extracted theta band power for inter-group comparison (Wide_RTBW contrast Narrow_RTBW). A significant negative cluster over prefrontal sensors (cluster-value = -32.543, p = 0.038; fig. 2c) was also found. Pearson’s correlation analysis between the theta power averaged over the significant cluster and RTBW revealed a significant negative correlation. Based on the above analysis results, we can conclude that the prestimulus WPE can predict the subjects’ ability in the temporal perception in visual leading SOA. And it is related to prestimulus theta band power.

**Fig. 2.**
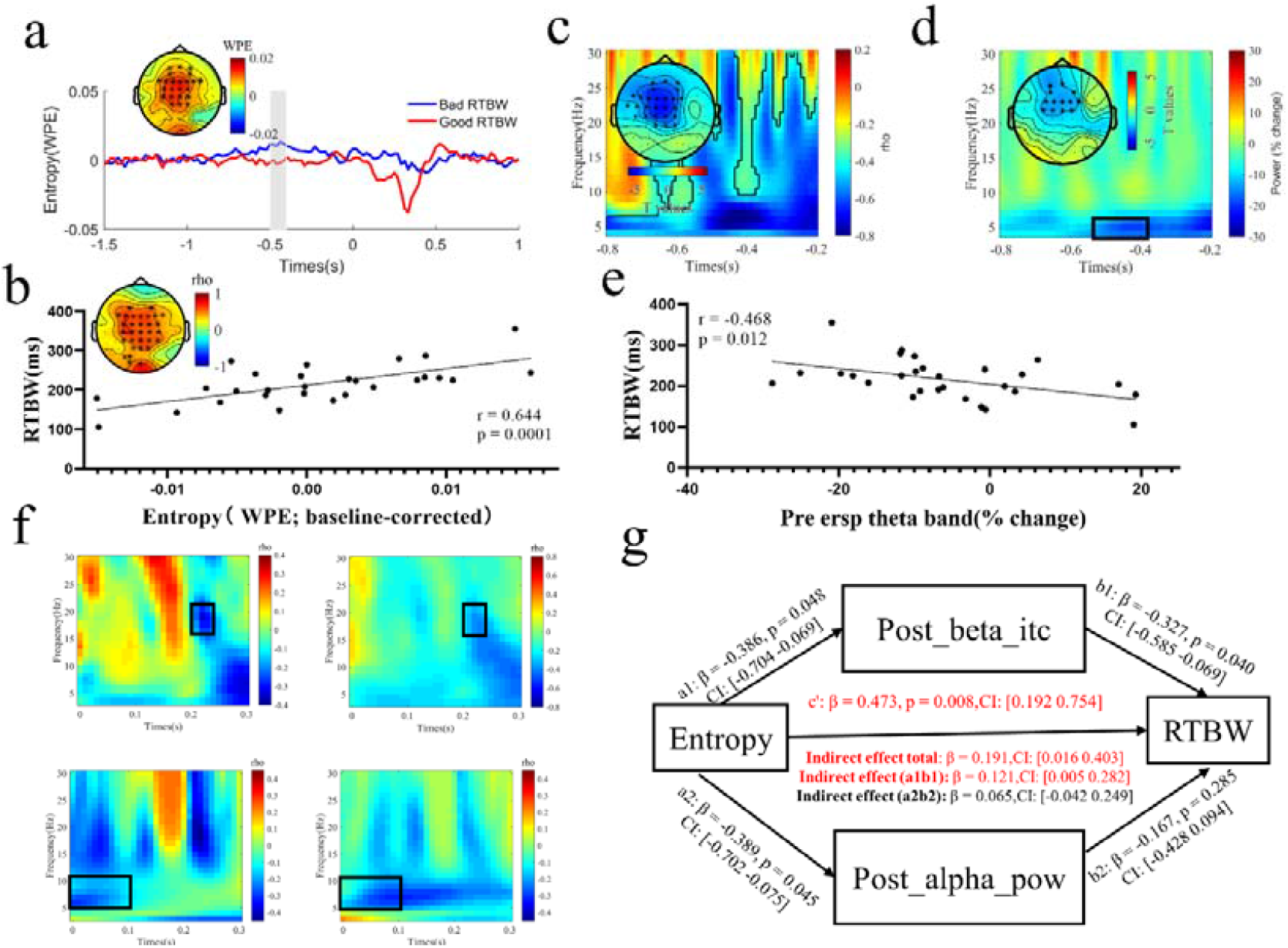
Inter-difference in the RTBW indexed by prestimulus WPE. (a) The time courses of average WPE over significant clusters for Wide_RTBW (blue line) and Narrow_RTBW (red line) groups are shown (baselined relative to −1.5 s to −1 s). Gray areas mark the temporal intervals (−0.5 to -0.4 s) with significant between-group differences. Inset is the topography over the significant time intervals where the color corresponds to the WPE difference between Wide_RTBW and Narrow_RTBW groups. Electrodes belonging to significant clusters are highlighted as asterisks. (b) Pearson’s correlation between the average prestimulus WPE over the significant cluster and RTBW. Colors on topography correspond to average correlation coefficients rho. (c and d) The time-frequency representation of prestimulus Pearson’s correlation between WPE and power in (c) and prestimulus power difference between Wide_RTBW and Narrow_RTBW groups in (d) averaged over significant channels, which are highlighted by asterisks in the inset topography. Black contours are the significant clusters. (e) Pearson’s correlation between the prestimulus theta band and right TBW. (f) The time-frequency representation of Pearson correlation coefficients over all 60 channels between prestimulus WPE and poststimulus ITC in the top-left subpanel; poststimulus ITC and right TBWs in the top-right subpanel; prestimulus WPE and poststimulus power in the bottom-left subpanel; poststimulus power and right TBWs in the bottom-right subpanel. The time-frequency point of interest is within the black rectangle area. (g) A parallel mediation model: poststimulus beta ITC partially mediated the relationship between prestimulus WPE and right TBWs. Red indicates that the path coefficient is significant.

Previous studies have shown that the prestimulus WPE is associated with poststimulus low-frequency power and ITC ^[13, 14]^. We constructed a parallel mediation model to examine whether prestimulus WPE influences perceptual response by shaping poststimulus-evoked activities. We first examined the Brain-Brain model (prestimulus WPE and poststimulus ITC/power) and the Brain-Behavior model (poststimulus ITC/power and RTBW) using Pearson’s correlation analysis within 0-0.3 s time range which was considered as the stage of sensory representation ^[23, 29]^ (fig. 2f). We found that in the time interval of 0.21 to 0.25 s, the frequency range of 16-20 Hz, and the sensors located in central and parietal regions ([C1, CZ, C2, CP1, CPZ, CP2, P1, PZ, P2], fig. 2f), there was a consistent negative correlation in the Brain-Brain model and Brain-Behavior model for ITC while the consistent negative correlation was found in the time interval of 0 to 0.1 s, the frequency range of 5-10 Hz for power. We then extracted and entered the data into a parallel mediation model. We controlled for potential confounds by including SOA, which is different across subjects, as covariates of no interest in the regression analysis. We found the correlation between prestimulus WPE and RTBW could be partially mediated by the poststimulus ITC but not power (indirect effect: a1b1 = 0.121, 90% CI [0.005 0.282]; a2b2 = 0.065, 90% CI [-0.042 0.249]; direct effect: c’= 0.473, 90% CI [0.192 0.754]; fig. 2g), suggesting higher prestimulus WPE might lead to worse temporal perception by partially leading to lower ITC which is associated with greater response time variability across trials ^[35]^.

We use the same analysis steps above to analyze the LTBW group. The prestimulus WPE averaged over -700 to -500 ms before the onset of the auditory stimulus was extracted and entered into an independent samples t-test (Wide_LTBW contrast Narrow_LTBW) combined with a spatial cluster-based permutation test. We found a significant positive cluster over left prefrontal sensors (cluster-value = 23.826, p = 0.046; fig. 3a), suggesting an increase in prestimulus WPE reflected weaker temporal perception. Positive correlations were significant between the prestimulus WPE and the magnitude of LTBW over the same left prefrontal sensors (cluster-value = 17.028, p = 0.046; fig. 3b). We found a negative cluster with a significant trend (cluster-value = -12729, p = 0.0959; fig. 3b) between the prestimulus WPE and power. A significant negative cluster over the central and parietal sensors (cluster-value = -35.667, p = 0.046; fig. 3b) was found when we limited the time range to -0.8 to -0.6s and the frequency range to a 20-30 Hz beta band. Again, a significant negative cluster (cluster-value = -43.232, p = 0.020; fig. 3c) over central and parietal sensors was found in between-group comparison (Wide_LTBW contrast Narrow_LTBW). The same significant negative correlation was found in Pearson’s correlation analysis between the beta power and LTBW. Based on the above analysis results, we can conclude that the prestimulus WPE can predict the subjects’ ability in the temporal perception in auditory leading SOA. Of note, prestimulus WPE is related to prestimulus beta band power but the distribution of significant brain regions is different.

**Fig. 3.**
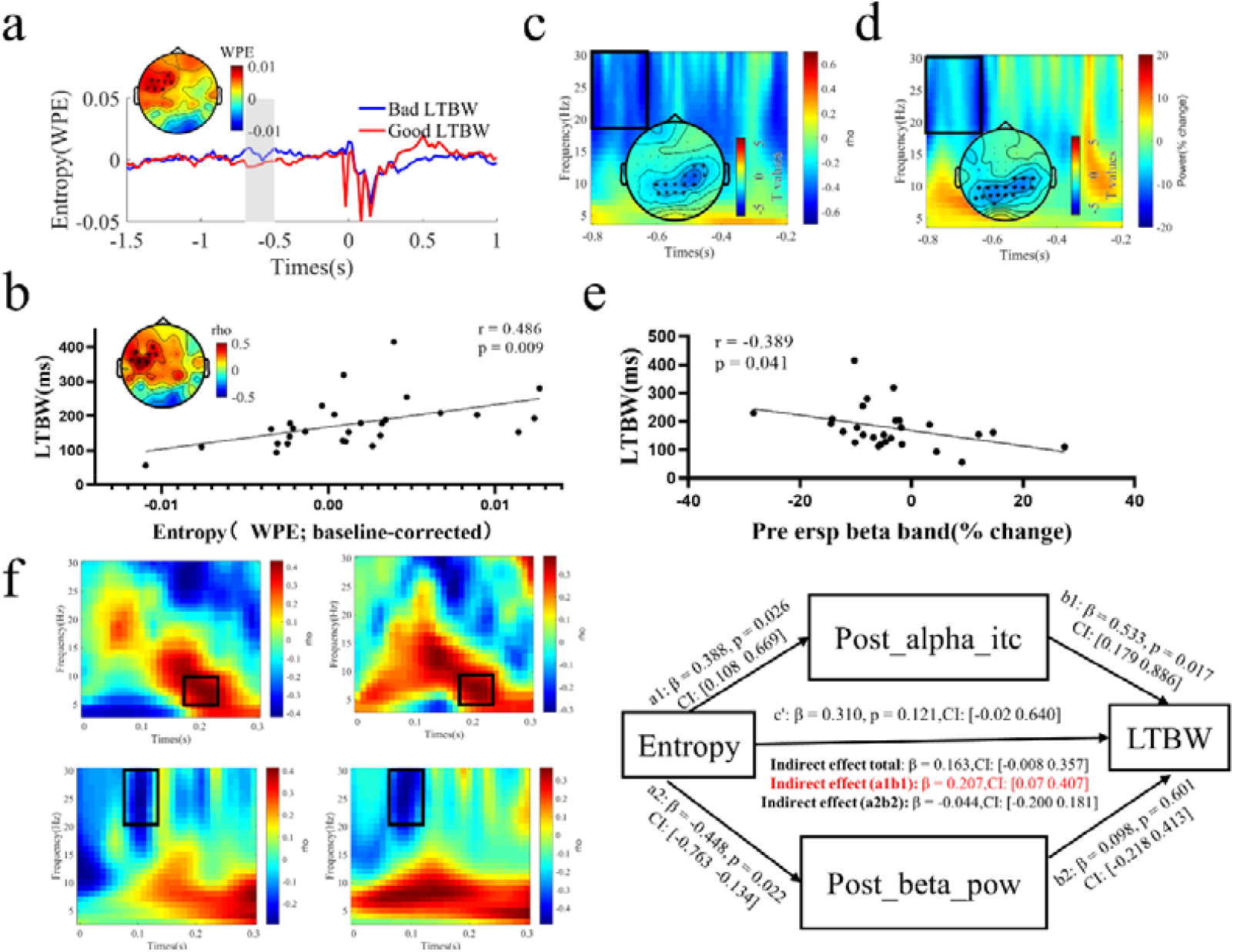
Inter-difference in the left TBW indexed by prestimulus WPE. (a) The time courses of average WPE over significant clusters for Wide_LTBW (blue line) and Narrow_LTBW (red line) groups are shown (baselined relative to −1.5 to −1 s). Gray areas mark the temporal intervals (−0.7 to -0.5 s) with significant between-group differences. Inset is the topography over the significant time intervals where the color corresponds to the WPE difference between Wide_LTBW and Narrow_LTBW group. Electrodes belonging to significant clusters are highlighted as asterisks. (b) The Pearson correlation between the average prestimulus WPE over the significant clusters and right TBW. Colors on topography correspond to average correlation coefficients r. (c) The time-frequency representation of prestimulus Pearson correlation between WPE and power averaged over all 60 channels. (e) The Pearson correlation between the prestimulus theta band and left TBW. (d) The time-frequency representation of prestimulus power difference (baseline-corrected relative to −1.5 to −1 s) averaged over significant channels which are highlighted by asterisks in the inset topography. The time-frequency point of interest is within the black rectangle area. (f) The time-frequency representation of Pearson correlation coefficients over all 60 channels between prestimulus WPE and poststimulus ITC in the top-left subpanel; poststimulus ITC and left TBWs in the top-right subpanel; prestimulus WPE and poststimulus power in the bottom-left subpanel; poststimulus power and right TBWs in the bottom-right subpanel. The time-frequency point of interest is within the black rectangle area. (g) A parallel mediation model: poststimulus alpha ITC fully mediated the relationship between prestimulus WPE and left TBWs. Red indicates that the path coefficient is significant.

We found that in the time interval of 0.19 to 0.24 s, the frequency range of 5-10 Hz, and the frontal, central, and parietal sensors ([FC1, FCZ, FC2, CP1, CPZ, CP2, C1, CZ, C2]; fig. 3f), there was a consistent positive correlation in the Brain-Brain model and Brain-Behavior model for ITC while the consistent negative correlation was found in the time interval of 0.08 to 0.13 s, the frequency range of 23-30 Hz involving power. So we extracted and averaged the raw data, and entered the data into a parallel mediation model. We included the SOA as covariates in the regression analysis. The results showed that the correlation between prestimulus WPE and LTBW could be fully mediated by the poststimulus alpha ITC but not power (indirect effect: a1b1 = 0.207, 90% CI [0.07 0.407]; a2b2 = -0.044, 90% CI [-0.2 0.181]; direct effect: c’= 0.310, 90% CI [-0.02 0.640]; fig. 3g), suggesting higher prestimulus WPE might lead to worse temporal perception by leading to higher ITC.

### 3.4 intra-individual differences indexed by prestimulus WPE mediated by prestimulus power and poststimulus ITC

Waschke et al. (2017) found higher neural irregularity quantified by prestimulus WPE was associated with optimized sensory encoding in auditory perception. We want to examine the relationship in the audiovisual domain.

In the visual leading SOA, we found no significant clusters. The cluster corresponding to the smallest p-value is negative (cluster-value = -25.441, p = 0.474; fig. 6a). However, a significant positive cluster was found between synchronous and asynchronous response within -0.2 to -0.1 s time interval (cluster-value = -218.979, p = 0.01; fig. 6b) over posterior sensors. We further found a significant negative Pearson correlation between prestimulus WPE and prestimulus power over all 60 channels and -1 to 0 s time interval (cluster-value = -54346, p = 0.002; fig. 6c).

Given that prestimulus WPE determined the perceptual response and was associated with prestimulus power, poststimulus jITC, and power, we aimed to examine whether prestimulus WPE influenced perceptual responses through prestimulus power as well as poststimulus jITC and power at the single-trial level with a parallel-serial mediation model. First, we tried to use linear or logistic regression models (behavioral response as dependent variable) to discover the consistent relationship between the above variables among subjects in statistics. A significant negative relationship between prestimulus WPE and perceptual response (synchronous responses are coded as 1, asynchronous responses are coded as 0) within the prestimulus -200 to -100 ms time window (cluster-value = -46.287, p = 0.004; fig. 6d) over posterior sensors. With increasing prestimulus WPE, the probability of asynchronous responses increased. Next, the prestimulus WPE averaged over P1 channels with the largest t-value in statistics was extracted as the predictor, and poststimulus jITC as the dependent variable, we found a significant positive relationship within the poststimulus 150 to 220 ms time window and 8-16 Hz frequency band (cluster-value = 56.758, p = 0.006; fig. 6e) over posterior sensors. However, we only found a marginally significant negative relationship between poststimulus jITC and perceptual response within the poststimulus 130 to 200 ms time window and 8-14 Hz frequency band (cluster-value = -18.762, p = 0.0699; fig. 6e) over the left parieto-occipital sensors. Further, we found a significant negative correlation between prestimulus WPE and poststimulus power across all time, frequencies, and channels (cluster-value = -379.751, p = 0.002; fig. 6e). A significant positive correlation between poststimulus power and perceptual response within the poststimulus 120 to 180 ms time window and 15-30 Hz frequency band (cluster-value = 28.948, p = 0.022; fig. 6e) over left parieto-occipital sensors was found (fig. 4e). Based on the relationship found between the above variables, we entered prestimulus WPE (averaged over -200 to -100 ms time window), prestimulus power (averaged over -200 to -100 ms time window and 9 to 14 Hz frequency band), poststimulus jITC (averaged over 140 to 200 ms time window and 9 to 14 Hz frequency band), poststimulus power (averaged over 120 to 180 ms time window and 15 to 30 Hz frequency band) averaged over the left parieto-occipital channel (PO3, PO5, PO7) of interest to a parallel-serial mediation model. We found the correlation between prestimulus WPE and perceptual response could be fully mediated by the prestimulus alpha power and poststimulus alpha ITC but not poststimulus beta power (indirect effect: a1b1 = -0.0013, 95% CI [-0.005 0.002]; a2b2 = -0.04, 95% CI [-0.009 0.0005]; a31b33 = -0.017, 95% CI [-0.037 0.003]; a31b32b1 = -0.001, 95% CI [-0.003 -0.0002]; a31b33b2 = -0.003, 95% CI [-0.007 0.0003]; direct effect: c’= -0.042, 95% CI [-0.106 0.022]; fig. 6f), suggesting higher prestimulus WPE might lead to better temporal perception by inducing lower alpha power which can cause higher ITC.

**Fig. 4.**
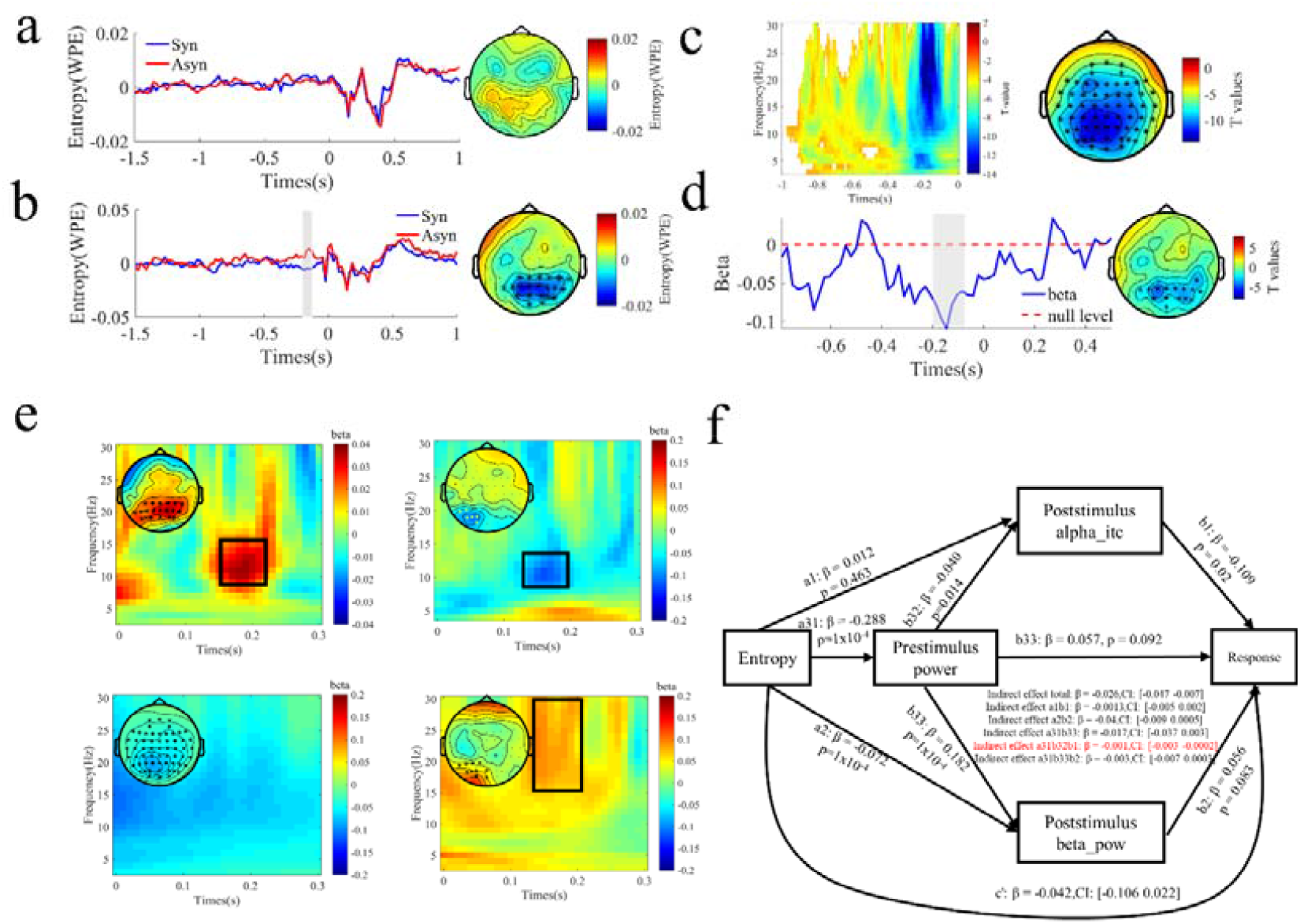
Intra-difference in the audio and visual leading SOA indexed by prestimulus WPE. (a) The time courses of average WPE over all 60 channels for synchronous (blue line) and asynchronous (red line) responses are shown (baselined relative to −1.5 to −1 s). The right panel is the topography over the -0.8 to -0.1 s time intervals where the color corresponds to the WPE difference between synchronous and asynchronous response. (b) The time courses of average WPE over significant cluster which is highlighted as asterisks in the right panel for synchronous (blue line) and asynchronous (red line) responses are shown (baselined relative to −1.5 to −1 s). Gray areas mark the temporal intervals (−0.2 to -0.1 s) with significant between-response differences. The right panel is the topography over the significant time intervals where the color corresponds to the WPE difference between synchronous and asynchronous response. (c) The time-frequency representation of prestimulus Pearson correlation between WPE and power averaged over the significant cluster marked by asterisks in the right topography. (d) The time course of beta regression coefficients (blue line) and null level regression coefficients averaged over the significant channels marked by asterisks in the right topography. (e) The time-frequency representation of regression coefficients averaged over significant channels marked by asterisks or marginally significant channels marked by dots between prestimulus WPE and poststimulus jITC in the top-left subpanel; poststimulus jITC and response in the top-right subpanel; prestimulus WPE and poststimulus power in the bottom-left subpanel; poststimulus power and responses in the bottom-right subpanel. The time-frequency point of interest is within the black rectangle area. (f) A parallel-serial mediation model: prestimulus alpha power and poststimulus alpha ITC fully mediated the relationship between prestimulus WPE and behavior responses. Red indicates that the path coefficient is significant.

## 4 Discussion

Prestimulus brain response directly influences perceptual decision-making or indirectly by task-related evoked activity ^[29]^. In this study, we use WPE, which reflects neural variability and is closely related to slow-wave activity amplitude, to test the above opinion at inter- and intra-individual levels. At the inter-individual level, prestimulus WPE predicts the width of individual TBWs mediated by poststimulus ITC, which might play a role in representing sensory evidence. At the intra-individual level, prestimulus WPE modulates the perceptual response by shaping the sensory representations indexed by poststimulus ITC at the single-trial level.

### 4.1 prestimulus WPE predicts the individual’s TBWs by shaping poststimulus sensory representation

A recent study found that prestimulus absolute WPE (nonbaseline-corrected) increased with age ^[48]^ and concluded that WPE is a trait-like marker of inter-individual difference. Our paper found that prestimulus relative WPE (−1.5 to -1 s baseline-corrected) is a state-like and task-dependent marker of inter-individual difference in young adults in audiovisual temporal perception. Since EEG entropy receives contributions from periodic and aperiodic signal parts ^[13, 14]^, we correlated it with prestimulus low-frequency relative power (−1.5 to -1 s baseline-corrected). Interestingly, we found prestimulus WPE is negatively associated with theta and high beta band power but not the alpha band power in visual-leading and auditory-leading SOA conditions. Several reasons might explain the results. First, most papers found that task-related not at rest or immediately before the task alpha power correlates with individual performance ^[16, 35]^. Second, the tasks in the studies that found spontaneous alpha power effect are self-related and complex ^[17, 49]^. Third, the subjects in our study are young adults with developing brains ^[50]^. Next, given the different roles of WPE in different SOAs, we will discuss them separately.

In visual-leading SOA, we found increasing prestimulus frontal WPE is associated with larger RTBW and decreasing theta power. Prestimulus frontal theta power is related to top-down expectation, attention, and preparation ^[51, 52]^. Furthermore, using an equiprobable Go/NoGo paradigm, a study found increasing prestimulus theta power was associated with early sensory processing enhancement ^[53]^. In addition, we also found poststimulus low beta ITC mediated the relationship between prestimulus frontal WPE and individual RTBWs. Non-human animal research indicated that low-beta (16–20 Hz) synchrony in the lateral prefrontal cortex (PFC) boosts the signal of sensory inputs ^[54]^. Combined with the above evidence, we speculated that prestimulus frontal WPE exerts a role in top-down mental readiness ^[51, 52, 55, 56]^, which could influence the evoked response related to sensory processes ^[53]^, then shape the magnitude of individual RTBWs.

In auditory-leading SOA, we found increasing prestimulus left-frontal WPE is associated with larger LTBW and decreasing central and parietal beta power. Moreover, the parallel mediation model indicated that the central theta/low-alpha ITC mediated the relationship between prestimulus left-frontal WPE and individual LTBWs. Studies related to temporal integration have shown that increased prestimulus parietal beta power is associated with larger cognitive (and attentional) control in the visual ^[57]^ and audiovisual domains ^[58]^. Furthermore, several previous studies indicated theta band ITC is related to auditory perceptual performance in humans ^[59, 60]^ and non-human monkeys ^[61]^. According to the topography and latency of theta/low-alpha ITC, we thought it was related to auditory sensory encoding and representation. So we concluded that prestimulus left-frontal WPE is related to cognitive control and can predict individual LTBWs by modulating poststimulus auditory representation.

### 4.2 prestimulus WPE shape temporal perception mediated by amodal representation at the intra-individual level

Our study found that prestimulus WPE shape individual perceptual judgments in auditory-leading SOA but not visual-leading SOA. Our results extended the findings of previous studies, which indicated that prestimulus alpha power affects audiovisual time perception judgment ^[48]^, showing that prestimulus WPE can also quantify the intra-individual difference in multisensory synchrony perception. Of note, the reason that prestimulus WPE had no role in perceptual judgments in visual-leading SOA might be that EEG entropy includes periodic and aperiodic signal components. So the non-periodic part of the entropy may interfere with the periodic one, resulting in its insufficient sensitivity as an indicator for quantifying intra-individual differences mainly modulated by the periodic power. Next, we performed several serial mediation models to test whether prestimulus brain states shape perceptual judgments by sensory representations marked by poststimulus-evoked activities ^[28, 29]^. The results showed that prestimulus WPE shape perceptual responses fully mediated by prestimulus alpha power and poststimulus alpha jITC in the parietal cortex at the single-trial level.

Many papers have shown that decreased prestimulus alpha power in the occipital-parietal areas is associated with enhanced neural excitability with visual ^[62]^ and audiovisual stimuli ^[48]^. So we thought the parietal WPE is also associated with neural excitability. As a sensory association region, the parietal cortex receives inputs from the auditory, visual, and somatosensory cortex, and so on. It is considered a multisensory integration region ^[63, 64]^. Walsh (2003 a,b) ^[65, 66]^ proposed a theory of magnitude (ATOM) that establishes commonalities in the sensory representations of various magnitudes, such as time, space, number, size, and speed, in the parietal cortex ^[67]^. Recent experimental studies have provided evidence supporting this theory ^[68, 69]^. Van Wassenhove (2009) also advised that time was in an amodal representational space in the parietal cortex. Taking the above evidence together, we thought that prestimulus WPE affects temporal perception mediated by prestimulus excitability and amodal representation in the parietal cortex.

It is worth noting that not all significant channels correlated with prestimulus WPE and behavioral response functioned in the mediation models. So the task-evoked response might explain the partial variance in mediating the prestimulus brain state and behavioral response.

## 5 conclusion

In this study, our objective was to investigate whether prestimulus neural variability, quantified using weighted permutation entropy (WPE), serves as an indicator of within-individual and between-individual variability in audiovisual temporal perception. Furthermore, we aimed to explore whether this variability has a direct impact on behavior or an indirect effect through post-stimulus-evoked responses. Our results revealed that prestimulus WPE effectively reflects intra-individual and inter-individual differences in audiovisual temporal perception. Specifically, an increase in prestimulus frontal WPE was found to predict narrower individual temporal binding windows (TBWs). The prestimulus frontal WPE played a top-down role in facilitating mental readiness for sensory processes in visual-leading SOA conditions, while it played a cognitive control role in auditory-leading SOA conditions. Additionally, prestimulus parietal WPE was associated with more asynchronous responses, indicating its role in indexing cortical excitability. Importantly, we observed that prestimulus WPE influences within-individual and between-individual variability in audiovisual temporal perception, mediated by sensory representations through mediation models.

## Acknowledgments

This work was supported in part by the National Key Research & Development Program of China (No. 2022YFF1202400) and the National Natural Science Foundation of China (No.62276181).

